# Comparative proteomics of octocoral and scleractinian skeletomes and the evolution of coral calcification

**DOI:** 10.1101/2019.12.30.891028

**Authors:** Nicola Conci, Martin Lehmann, Sergio Vargas, Gert Wörheide

**Affiliations:** Department of Earth and Environmental Sciences, Ludwig-Maximilians-Universität, Munich, Germany; Biozentrum der LMU München, Department of Biology I–Botany, Grosshaderner Strasse 2–4, Planegg-Martinsried, Germany; SNSB - Bayerische Staatssammlung für Paläontologie und Geologie, Munich, Germany; GeoBio-Center LMU, Ludwig-Maximilians-Universität, Munich, Germany

## Abstract

Corals are ecosystem engineers of the coral reefs, one of the most biodiverse but severely threatened marine ecosystems. The ability of corals to form the three dimensional structure of reefs depends on the precipitation of calcium carbonate under biologically control. However, the exact mechanisms underlying this biologically controlled biomineralization remain to be fully unelucidated, for example whether corals employ a different molecular machinery for the deposition of different calcium carbonate (CaCO_3_) polymorphs (i.e., aragonite or calcite). Here we used tandem mass spectrometry (MS/MS) to compare skeletogenic proteins, i.e., the proteins occluded in the skeleton of three octocoral and one scleractinian species: *Tubipora musica* and *Sinularia* cf. *cruciata*, both forming calcite sclerites, the blue coral *Heliopora coerulea* with an aragonitic rigid skeleton, and the scleractinian aragonitic *Montipora digitata*. We observed extremely low overlap between aragonitic and calcitic species, while a core set of proteins is shared between octocorals producing calcite sclerites. However, the same carbonic anhydrase (CruCA4) is employed for the formation of skeletons of both polymorphs. Similarities could also be observed between octocorals and scleractinians, including the presence of a galaxin-like protein. Additionally, as in scleractinians, some octocoral skeletogenic proteins, such as acidic proteins and scleritin, appear to have been secondarily co-opted for calcification and likely derive from proteins playing different extracellular functions. In *H. coerulea*, co-option was characterized by aspartic acid-enrichment of proteins. This work represents the first attempt to identify the molecular basis underlying coral skeleton polymorph diversity, providing several new research targets and enabling both future functional and evolutionary studies aimed at elucidating the origin and evolution of biomineralization in corals.

## Introduction

The capacity of animals to actively control the deposition of mineral skeletons has been a long debated topic, with different models of calcification being proposed over the years. In the ‘*organic matrix mediated*’ (Lowenstam 1981) or ‘*biologically controlled*’ (Mann 1983) scenario, an animal employs sets of macromolecules to guide the deposition of its mineral skeletal structures. In line with this, several biomineralization-related processes including crystal nucleation and growth (Liu et al. 2012; Wheeler et al. 1981; Mitterer 1978; Von Euw et al. 2017), or the induction of a given calcium carbonate (CaCO_3_) polymorph (i.e. aragonite and calcite) (Amos et al. 2010; Falini et al. 1996; Goffredo et al. 2011; Rahman et al. 2011) appear to be regulated by proteins included in the skeleton organic matrix (OM): a diverse array of proteins, polysaccharides (Goldberg 2001; Naggi et al. 2018), and lipids (Farre & Dauphin 2009; Farre et al. 2010) occluded within the mineral fraction of the skeleton;. Over the last few years, advances in proteomic research have enabled the simultaneous characterizations of several OM proteins in different groups of marine calcifying invertebrates, including molluscs (Marie et al. 2010, 2013; Mann & Jackson 2014), corals (Drake et al. 2013; Ramos-Silva et al. 2013; Takeuchi et al. 2016), brachiopods (Jackson et al. 2015) and echinoderms (Mann et al. 2008; Flores & Livingston 2017; Flores et al. 2016). These studies showed that invertebrate skeletal proteomes include varying fractions of novel proteins - producing no significant matches against DNA sequence databases - and do exhibit contrasting rates of conservation between and within lineages. For instance, about 40% of the skeletal proteome is shared among echinoderms (Flores & Livingston 2017), while in molluscs the fraction of shared proteins is only about 10% (Kocot et al. 2016). Although a core set of proteins appears to be conserved across molluscs, irrespective of the morphological features of the shell (Arivalagan et al. 2017), the occurrence of both aragonite and calcite layers within the shell allowed Marie et al. (2012) to compare skeleton organic matrix proteins (SOMPs) associated with different CaCO_3_ polymorphs. The different proteins specifically associated with the aragonitic or calcitic shell layers suggest that molluscs may use different molecular mechanisms for the deposition of these structures (Marie et al. 2012). In corals (class Anthozoa, phylum Cnidaria), putative relationships between structural characteristics of the skeleton, like its CaCO_3_ polymorph, and the molecular machinery employed for its formation, have hitherto only been marginally addressed, although some skeletogenic coral proteins have been suggested to drive the *in vitro* crystallization of specific calcium carbonate polymorphs (Goffredo et al. 2011; Rahman et al. 2011). However, no study to date has leveraged mass spectrometry-based protein discovery methods to characterize and compare skeletal proteomes across corals that exhibit different biomineralization strategies, i.e., those that produce aragonite vs. those that produce calcite. CaCO_3_ skeleton-producing anthozoan corals belong in two different clades, namely the order Scleractinia (stony corals; subclass Hexacorallia) and in the subclass Octocorallia (soft corals). As major contributors to CaCO_3_ deposition, scleractinians have been the focus of extensive biomineralization-related research, and skeletogenic proteomes have been characterized for different scleractinian species (Drake et al. 2013; Ramos-Silva et al. 2013; Takeuchi et al. 2016). However, the uniformity in biomineralization strategies (i.e., aragonitic exoskeleton) present among scleractinians makes this group inappropriate to investigate the biological regulation of skeletal polymorph deposition and its evolution. On the contrary, the occurrence within Octocorallia of both calcite and aragonite skeletons offers a unique opportunity to compare the skeletogenic repertoires associated with different skeletal structures and CaCO_3_ polymorphs. Despite this, information on octocoral biomineralization-related proteins is extremely limited (but see Debreuil et al. (2012; 2011) and Rahman et al. (2011)), and transcriptomic-proteomic coupled data is hitherto not available for this group.

Here, we used tandem mass spectrometry (MS/MS) to characterize the skeletogenic proteome of three soft coral species exhibiting different skeleton morphologies and mineralogies: the leather coral *Sinularia* cf. *cruciata* and the pipe organ coral *Tubipora musica*, both characterized by the production of calcite sclerites, and the massive, aragonitic blue coral *Heliopora coerulea*. To compare skeletogenic repertoires between scleractinians and aragonitic octocorals, we additionally examined the proteome of the stony coral *Montipora digitata.* Our work represents the first study of skeletome diversity across and within anthozoan corals providing 1) the identification of several new coral biomineralization-related proteins, and 2) a comparative analysis examining putative relations between polymorph and type of skeletal structure, and the molecular machinery employed by corals for its formation.

## Material and Methods

### Extraction of OM proteins

Samples of *T. musica*, *H. coerulea, Sinularia* cf. *cruciata* and *M. digitata*, cultured in research aquaria (closed artificial seawater systems) were bleached in 5% NaOCl (Sigma-Aldrich) for 72 hours to remove the tissue and other potential contaminants. They were subsequently rinsed several times with ultrapure water and oven-dried at 37°C. Clean skeletons were ground to powder with a mortar and pestle, and again bleached (5% NaClO solution for 5 hours), washed with ultrapure water and oven-dried at 37°C. The skeleton powder was decalcified with 10% acetic acid for 24 hours at room temperature on an orbital shaker. The decalcification solution was centrifuged (14,000 g, 30 min, 25°C) to separate the acid soluble (ASM) and insoluble (AIM) fractions. The obtained insoluble pellets were washed several times with ultrapure water, dried and stored at −80°C until further analysis. The supernatants (ASM) were desalted and concentrated using Amicon Ultrafiltration devices (15 ml, 3 kDa cut-off), and the ASM proteins were precipitated following the method described in Wessel and Flügge (1984). Briefly, four volumes of methanol, one of chloroform and three volumes of water were added to one volume of sample and the solution was centrifuged at 14,000 g for 20 min at 25°C. After centrifugation, the supernatant was discarded. Three volumes of methanol were added and the solution was centrifuged again. The resulting protein pellets were air-dried and stored at −80°C. For each species, two skeleton samples from the same colony were independently processed.

### SDS-PAGE Analysis

ASM and AIM proteins were dissolved in 2X Laemmli buffer (95% buffer - 5% beta-mercaptoethanol) (BioRad). As observed by Ramos-Silva et al. (2013), AIM pellets were only partly dissolved. Samples were denatured for 2 minutes at 95 °C and loaded on 12.5% polyacrylamide gels. SDS-PAGE was run on a BioRad MiniProtean Tetra Cell at constant voltage for ca. 70 minutes. Proteins were visualized after staining with ProteoSilver Silver Stain Kit (Sigma-Aldrich) (S.Fig.1) with the Precision Plus Protein™ Dual Color Standards (Biorad) as a size marker.

### Proteomic analysis

For mass spectrometry analysis, sample aliquots were loaded on a 12.5% acrylamide gel and run as described above. In an effort to reduce potential variability due to technical factors, we included replicates within the experimental design. These include triplicates for each OM fraction (soluble and insoluble) for each of the two per-species extractions. However, the presence of technical causes underlying the non-detection of SOMPs cannot be completely excluded. Three replicates per fraction per sample (n=6 per sample, n=12 per species) were excised from the gel and digested with trypsin prior to analysis on a Bruker Impact II Q-Tof mass spectrometer (Bruker Corp. Billerica, Massachusetts, USA) coupled with an Ultimate 3000 RSLC nano liquid chromatography (Thermo Fisher, Waltham, Massachusetts, USA). Peptide separation was performed using an Acclaim PepMap RSLC column with 75 μm diameter, 25 cm length, C18 particles of 2 μm diameter and 100 Å pore size (Thermo Fisher, Waltham, Massachusetts, USA). Data were analyzed using MaxQuant 1.5.2.8 (Cox & Mann 2008). Common contaminants, potential symbiont and bacterial sequences were filtered and peptides were mapped against sequence datasets for the target species. Sample processing and mass spectrometry were performed by the MSBioLMU Unit at the Biology Department I of the Ludwig-Maximilians University in Munich (Germany). Sequences with at least two unique matching peptides were considered for downstream analysis.

### Bioinformatic analysis of OM proteins

Identified OM proteins were annotated by Blastp (cut off e-value: 1e^−10^) against the NCBI non-redundant database. Distribution of homologs (S.Mat 1) of octocoral and *M. digitata* SOMPs was subsequently assessed within a set of cnidarian genomes and transcriptomes (Voolstra et al. 2015; Shinzato et al. 2011; Voolstra et al. 2017; Pratlong et al. 2015; Jeon et al.; Liew et al. 2016) with Blastp applying the same search criteria described above. Presence of signal peptide, transmembrane regions, GPI-anchor was predicted with SignalP 4.0 (Petersen et al. 2011), TMHMM 2.0 (Krogh et al. 2001) and PredGPI (Pierleoni et al. 2008) respectively. Protein isoelectric point was determined with ProtParam (Gasteiger et al. 2005). The amino acid composition of the acidic proteins detected was computed with a custom script available at https://gitlab.lrz.de/palmuc/Concietal_proteomics_skeletomes. Relative amino acid frequencies and distribution of aspartate residues within acidic proteins was determined on sequences predicted as complete after removal of the signal peptide sequence. For the distribution of aspartate residues, a frequency table of the distance between aspartate residues within a protein was first computed. Median distance values and amino acid frequencies were then used to perform principal component analysis (PCA) of acidic proteins.

### Phylogenetic Analyses

For phylogenetic inference, protein queries were blasted against a database of cnidarian sequences. The following e-value cutoffs were used: 1e^−05^ (scleritin), 1e^−20^ (carbonic anhydrase) and 1e^−50^ (hephaestin-like). Sequences predicted as ‘internal’ (i.e. lacking both 3’ and 5’ ends) by TransDecoder were discarded and a minimum length filter of 250 and 600 residues was applied to significant hits for carbonic anhydrase (CA) and hephaestin-like, respectively. For the former, sponge and human sequences used in Voigt et al. (2014) were added to the dataset. Carbonic anhydrases from the green algae *Chlamydomonas reinhaardtii* (P20507) and *Desmodesmus* sp. (AOL92959.1) were used as outgroup. Sequences were aligned with both MUSCLE (Edgar 2004) and MAFFT (Katoh & Standley 2013), and best-fit models were estimated with Prottest 3.4 (Darriba et al. 2011). Maximum-Likelihood Analysis was performed in Seaview 4 (Gouy et al. 2010) using PhyML 3.1 (Guindon & Gascuel 2003), while MrBayes 3.2 was used for bayesian inferences. Trees were sampled every 100^th^ generation (nruns=2) and burn-in fraction for each analysis was determined after visual inspection of the trace files using Trace v1.6 (available at http://tree.bio.ed.ac.uk/software/tracer). All alignments, trees, and protein sequences used for phylogenetic analyses are available at https://gitlab.lrz.de/palmuc/Concietal_proteomics_skeletomes

## Results

### Shared and species-specific components of the anthozoan skeletome

The discovery and subsequent annotation of anthozoan skeleton organic matrix proteins (SOMPs) retrieved between 12 and 54 proteins (S.Mat 1) with low protein numbers shared between octocoral species. However, simultaneously small sets of skeletogenic proteins shared between organisms at different taxonomic levels were identified. These included instances of proteins being secreted in the skeleton of both scleractinians and octocorals. The aragonitic octocoral *H. coerulea* did not exhibit higher similarity to aragonitic scleractinians compared to calcitic soft corals, suggesting that the CaCO_3_ polymorph had no noticeable effect on skeletome conservation between groups.

**Fig. 1.**
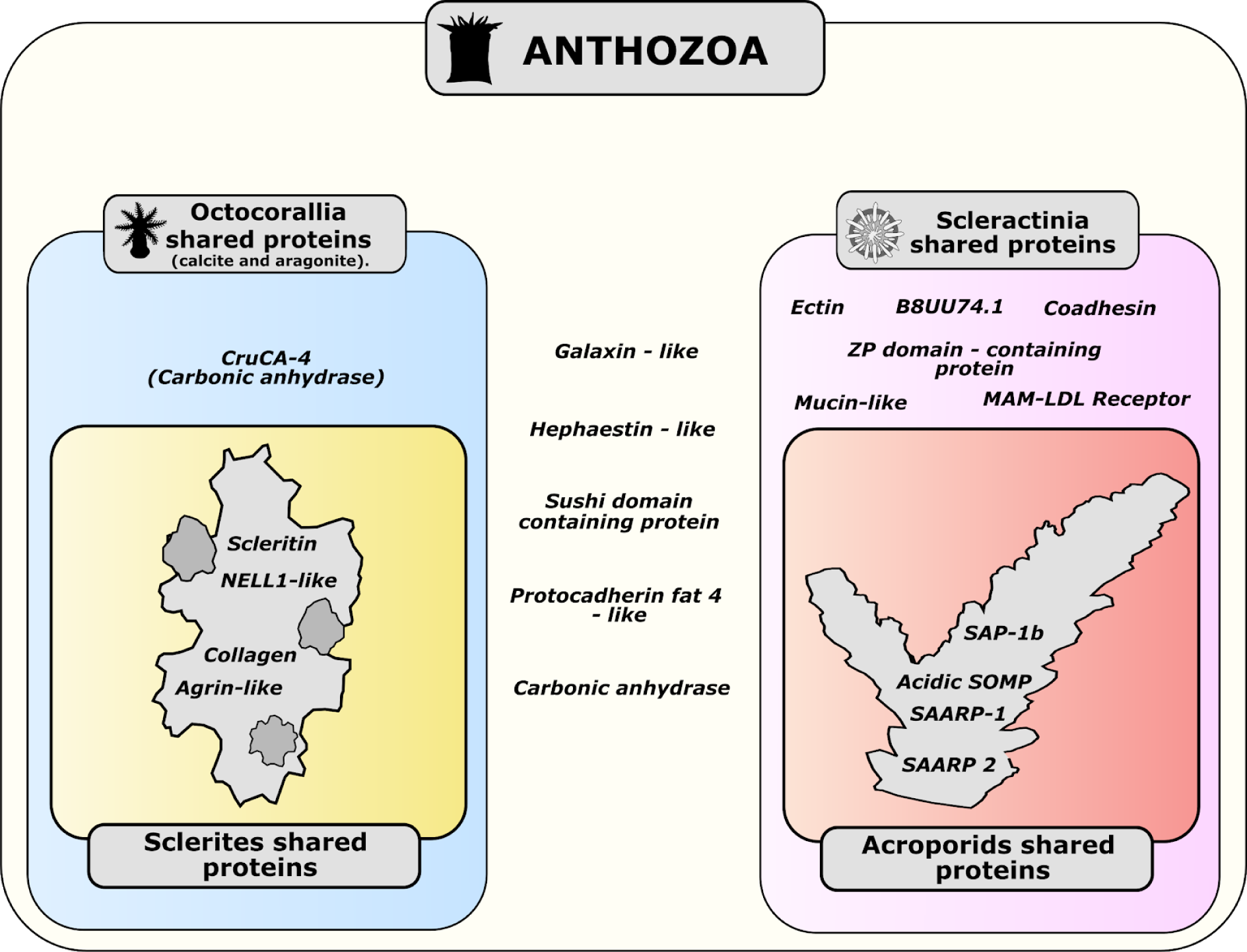
Overview of shared *skeletome* proteins across Anthozoa. Alcyonacea includes the proteomes of *T. musica* (this study) and *S.* cf. *cruciata* (this study). Calcitic octocorals: Alcyonacea + involvement of scleritin in *C. rubrum* (Debreuil et al. (2012)). Octocorallia: Alcyonacea + *H. coerulea* (this study) + involvement of CruCA4 in *C. rubrum* (Le Goff et al. (2016)). Acroporiidae: proteomes of *M. digitata* (this study) + *A. digitifera* (Takeuchi et al. (2016) + *A. millepora* (Ramos-Silva et al. (2013). Scleractinia: Acroporidae + proteome of *S. pistillata* (Drake et al. 2013).

Common to all four species analyzed is a Sushi-domain containing protein also containing NIDO (IPR003886), AMOP (IPR005533) and Von Willebrand factor D (IPR001846) domains. This same arrangement is present in the mucin-like B3EWY9 found in the *A. millepora* SOMP, and the proteins identified in this study also share local similarity with protein P13 from the skeleton of *S. pistillata.* Other proteins secreted in both scleractinian and octocoral skeletons include a putative homolog of galaxin, a hephaestin-like multicopper oxidase (MCO), one cell adhesion protocadherin Fat 4-like protein, and carbonic anhydrases. The first one, galaxin, was detected in the sclerites of *T. musica* (TR44621|c0_g1_i1) and we named it octogalaxin-1. In addition to the characteristic presence of multiple di-cysteine motifs, this protein is - as in scleractinians - predicted to be secreted and the signal peptide is followed by a R-X-R-R endoprotease target motif (Fukuda et al. 2003). The multicopper oxidase was found in *S.* cf. *cruciata* (TR42435|c0_g1_i1) and the protocadherin-like proteins were identified in the skeleton of *H. coerulea* (DN66065_c0_g1_i4). Although our comparative analysis of octocoral skeletal proteomes did not find evidence of a conserved *octocoral biomineralization toolkit*, we found homologs of *Corallium rubrum*’s carbonic anhydrase CruCA-4 (Le Goff et al. 2016) in both aragonitic and calcitic species. One CruCA4 homolog was found in *S. cruciata*, while in the aragonitic blue coral two homologs of CruCA4 were detected. As reported for *C. rubrum* (Le Goff et al. 2016; Del Prete et al. 2017), the histidine residue involved in the proton transfer is not conserved in all other homologs of the protein (S.Fig.2). Mutation of the His64 residues have been linked to decreases in efficiency (Vullo et al. 2008). Additionally, one of the two *H. coerulea* carbonic anhydrases (DN64689_c5_g1_i4) is predicted to be an acatalytic carbonic anhydrase-related protein (S.Fig.2). Four additional proteins are present in the sclerites of both calcitic octocorals analyzed and they represent best reciprocal hits between the two species. This sclerite “*toolkit”* includes 1) scleritin, 2) an agrin-like protein consisting of repeated Kazal domains (IPR036058) and one C-terminal WAP (IPR008197) domain, 3) a kinase C-binding, NELL1-like protein and 4) one collagen alpha-chain like protein. Two scleritin-like sequences (TR40200|c16_g1_i1 and TR42410|c0_g2_i1) were detected in *T. musica*, while only one match was produced in *Sinularia.* Agrin is a glycoprotein which in humans participates in cell-matrix interactions (Groffen et al. 1998), and agrin-like and agrin-like protease inhibitors have been recently found in the skeleton of the seastar *Patiria miniata* (Flores & Livingston 2017). NELL-1 is, on the other hand, involved in bone formation in vertebrates (Aghaloo et al. 2007; Zou et al. 2011). The NELL1-like protein identified here also exhibit local similarity to P32 (kielin-like), a secreted protein found in the skeleton of *S. pistillata* (Drake et al. 2013).

In the scleractinian *M. digitata*, with the exception of the secreted acidic protein SAP-1a, we retrieved the entire acidic protein repertoire previously isolated from the skeletons of *A. millepora* and *A. digitifera* (Ramos-Silva et al. 2013; Takeuchi et al. 2016). This includes both secreted aspartic acid-rich proteins SAARP-1 and SAARP-2, the acidic secreted organic matrix protein B3EWY7 and the secreted acidic protein SAP-1b. In addition, putative orthologs for *A. millepora* mucin-like (B3EWY9), coadhesin-like (B3EWZ3.1) and carbonic anhydrase (B8V7P3.1) were also present. Of note is the presence in *M. digitata* of a lithostatine-like protein containing a c-type lectin domain. The presence of this domain is a common feature for skeletogenic proteins in several marine invertebrates, such as mollusks (Mann et al. 2000; Matsubara et al. 2008; Weiss et al. 2000), see Sarashima et al. (2006) for a review), and birds (Mann & Siedler 2004, 2006). SOMPs with sequence similarity to lithostatin and c-type lectin-like proteins have been characterized from the skeletons of echinoderms (Wilt 2002) but have to our knowledge not been reported in corals to date.

### Similarity between scleractinian and octocoral acidic proteins

Two acidic proteins (DN60904_c0_g1_i1 and DN65627_c8_g3_i2) were detected in the organic matrix of *H. coerulea* and one in *T. musica* (TR43768|c0_g2_i1). Both do not currently match any published sequence available in public databases (Blastp e-value cut-off: 1e^−05^) outside of Octocorallia. As for scleractinians acidic SOMPs, homologs of *H. coerulea* acidic protein 1 exhibit higher isoelectric points which are related to lower aspartic acid contents. In an effort to investigate sequence similarities between octocoral and scleractinian acidic SOMPs and their non-acidic homologs in a phylogenetic independent way, we conducted a PCA analysis based on sequence amino acid composition and the distribution (i.e. median distance) of aspartate residues along the sequence (Fig. 2)

**Fig. 2.**
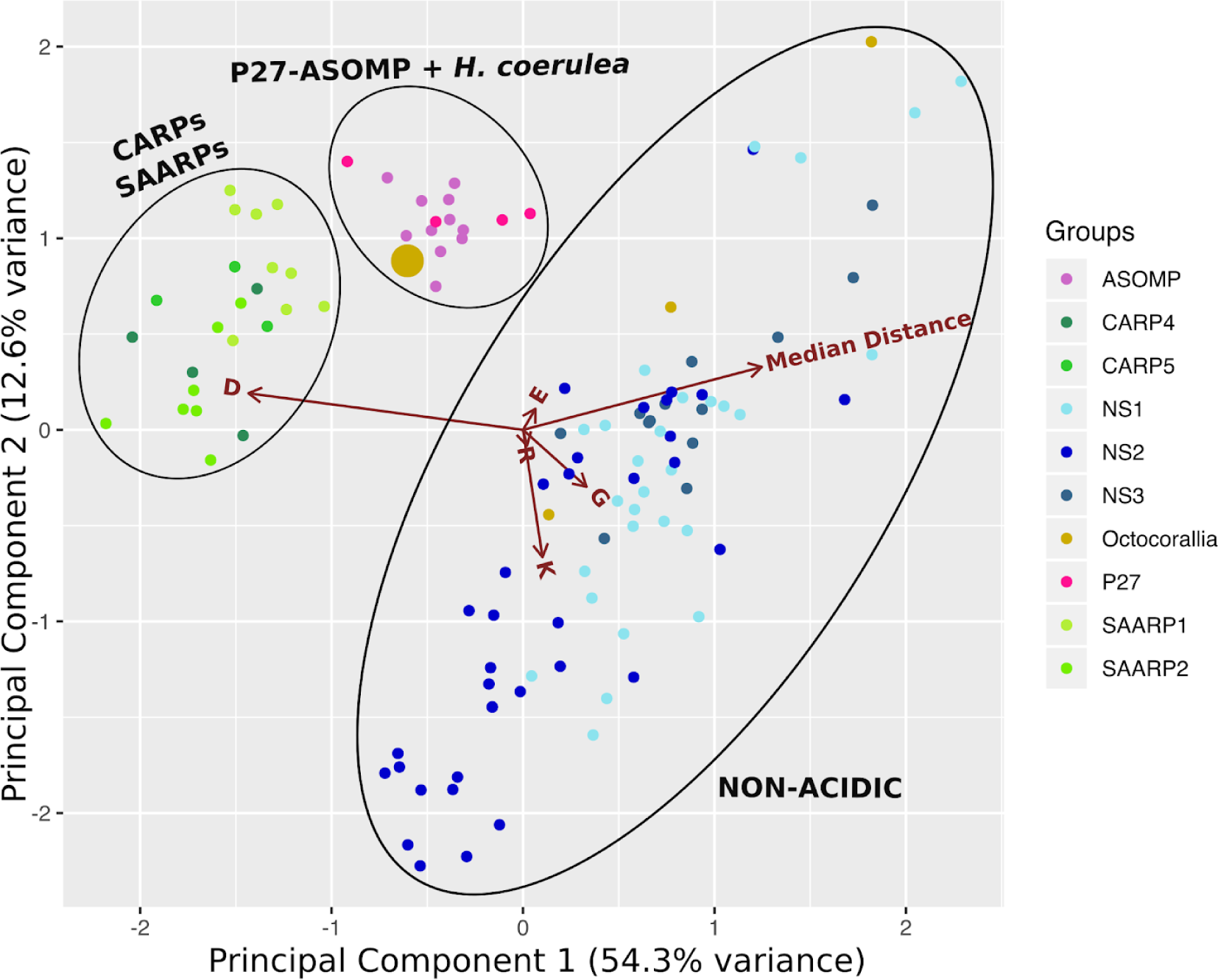
Principal component analysis (PCA) of anthozoan acidic SOMPs and their non-acidic homologs. PCA based on protein amino-acid composition and distribution - as median distance between residues - of aspartic acid along each sequence. Numbering of non-skeletogenic (NS) groups based on Conci et al. (2019). Only the 5 most contributing variables are displayed. Large golden circle: *H. coerulea* acidic protein-1 (DN60904_c0_g1_i1). Small golden circles: putative non-acidic homologs of *H. coerulea* acidic protein-1.

Despite not displaying significant similarity to scleractinian sequences, *H.coerulea* acidic protein-1 grouped together with the scleractinian acidic protein ASOMP-P27 (Ramos-Silva et al. 2013; Drake et al. 2013; Conci et al. 2019). Main sequence features contributing to the clustering patterns observed are similarities in aspartic acid content and distribution within the protein sequence. Non-acidic homologs of *H. coerulea* acidic protein-1 clustered with other non-acidic proteins. Apart from exhibiting lower aspartate contents, proteins within this group appear characterized by a higher lysine and glycine content compared to their acidic putative homologs.

### Evolutionary history of octocoral and scleractinian SOMPs

To further explore the evolutionary history of octocoral and scleractinian SOMPs we conducted phylogenetic analyses of protein sequences derived from the skeletome for scleritin, multicopper oxidases and carbonic anhydrases. Information on scleritin secretion into octocoral skeletons was integrated with previously estimated scleritin presence-absence data (Fig. 3a). Phylogenetic analysis split scleritin homologs into two distinct and well supported clades (Fig.3b). The three sequences identified in *T. musica* and *S.* cf. *cruciata* grouped together with the scleritin originally described in *C. rubrum* by Debreuil et al. (2012), alongside all other scleritin homologs found in octocoral species characterized by the presence of calcitic sclerites (Fig.3a).

**Fig. 3.**
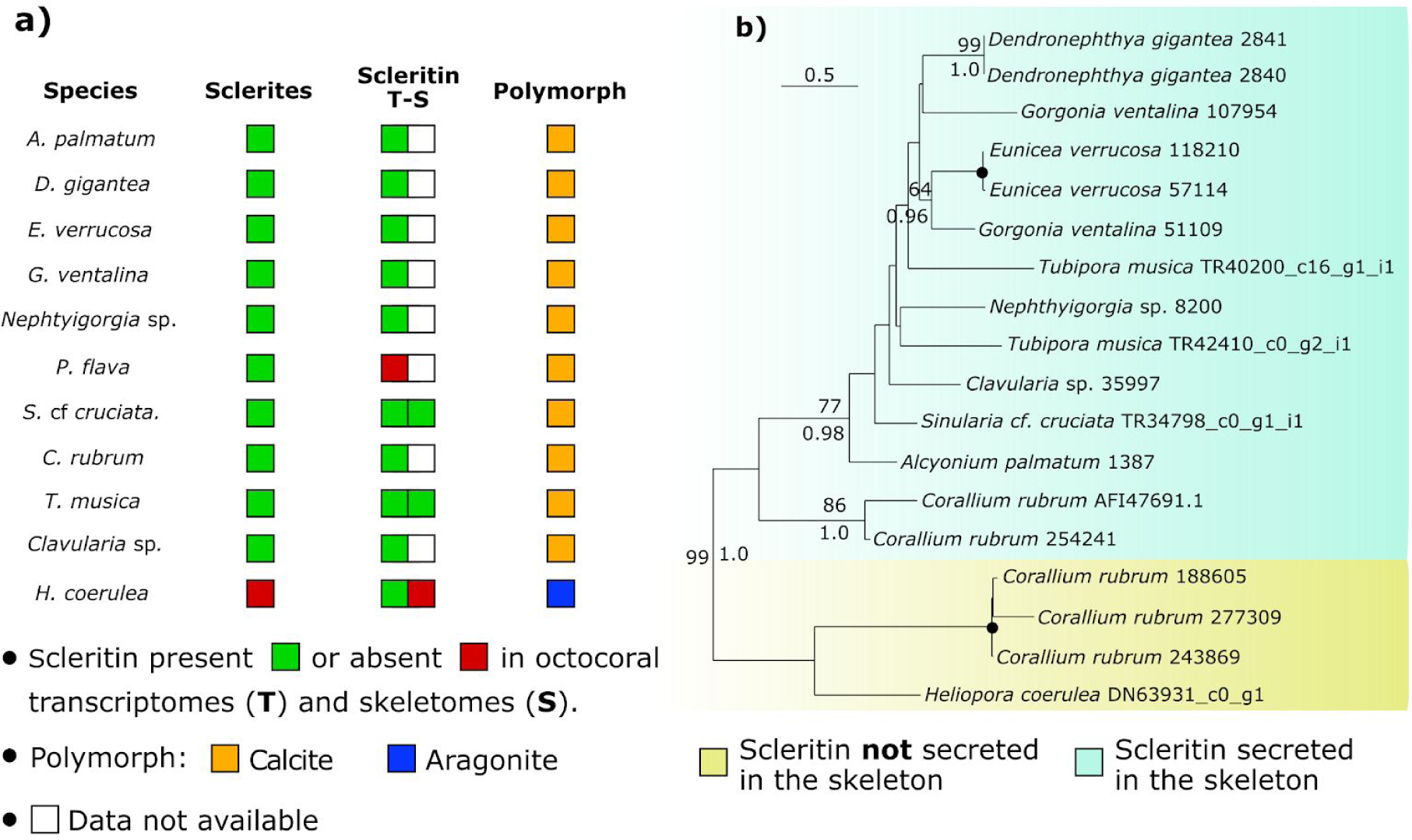
**a)** Presence-absence of scleritin in octocoral skeletomes in relation to skeletal structures. **b)** Phylogenetic analysis of scleritin. Protein sequences were aligned with MUSCLE and Maximum-Likelihood analysis (400 replicated) was done with Seaview 4. Bayesian analysis was performed with MrBayes 3.2. Black dot on node indicates full support (100% bootstrap value and 1.0 posterior probability). Involvement of scleritin in *C. rubrum* biomineralization based on (Debreuil et al. 2012). Phylogeny based on MAFFT aligning algorithm in S.Fig.3. All alignments available in S.Mat 2

We, therefore, referred to this clade as ‘skeletogenic’ since all the scleritin sequences implicated to date in octocoral biomineralization are comprised within it. A second group, termed ‘non-skeletogenic’ includes the scleritin-like protein expressed in the tissues but not found occluded in the skeleton of *H. coerulea*, and three other putative scleritin homologs found in *C. rubrum*.

To infer a phylogenetic tree for hephaestin-like proteins, putative homologs were searched across Cnidaria using the three multicopper oxidases described in Takeuchi et al (2016) as query. Each query protein formed a different clade populated by scleractinian and corallimorph sequences. Proteins present in scleractinian skeletons all grouped within clade 1.

**Fig. 4.**
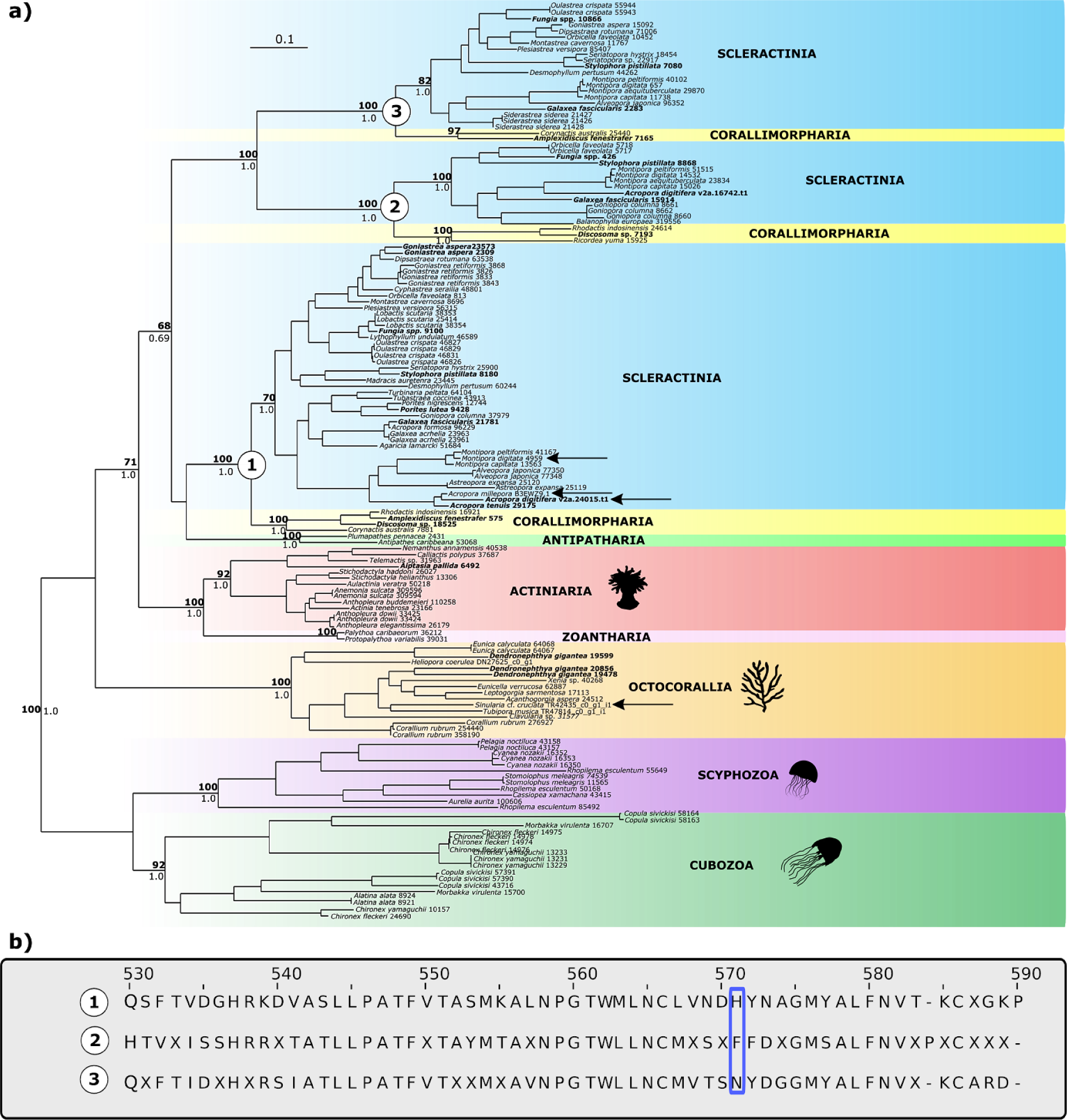
**a)** Phylogenetic analysis (400 bootstrap replicates) of cnidarian multicopper oxidases (MCOs). Aligning algorithm: MAFFT. Best-fit model: WAG+G+I. Number on nodes: bootstrap support and posterior probability values. Support values in bold: node supported (> 50) in MUSCLE-based phylogeny (S.Fig.4). **b)** Section of multiple consensus (60%) sequences alignment for the three ‘corallimorph + scleractinian’ clades. Blue box highlights the absence of the type-I copper binding histidine in clades 2 and 3. Histidine classification based on Takeuchi et al. (2016). All alignments available in S.Mat 2

All other cnidarian taxa formed well-supported monophyletic groups. Homologs identified in black corals (Antipatharia) grouped within clade 1 but with low support. Analysis of the consensus sequence alignment shows that one of the histidines involved in copper binding is not present in hephaestin-like proteins from clade 2 and 3, while all copper-binding residues listed in Takeuchi et al (2016) are conserved across clade 1 and all other cnidarian groups, including octocorals and the protein secreted in the sclerites of *S.* cf. *cruciata*.

Finally, all homologs of the carbonic anhydrase CruCA4 occupied the same clade (Fig. 5). This group also included the *H. coerulea* CA-related protein we found in the skeleton of this species. Scleractinian biomineralization related CAs did, on the other hand, split into three distinct groups.

**Fig. 5.**
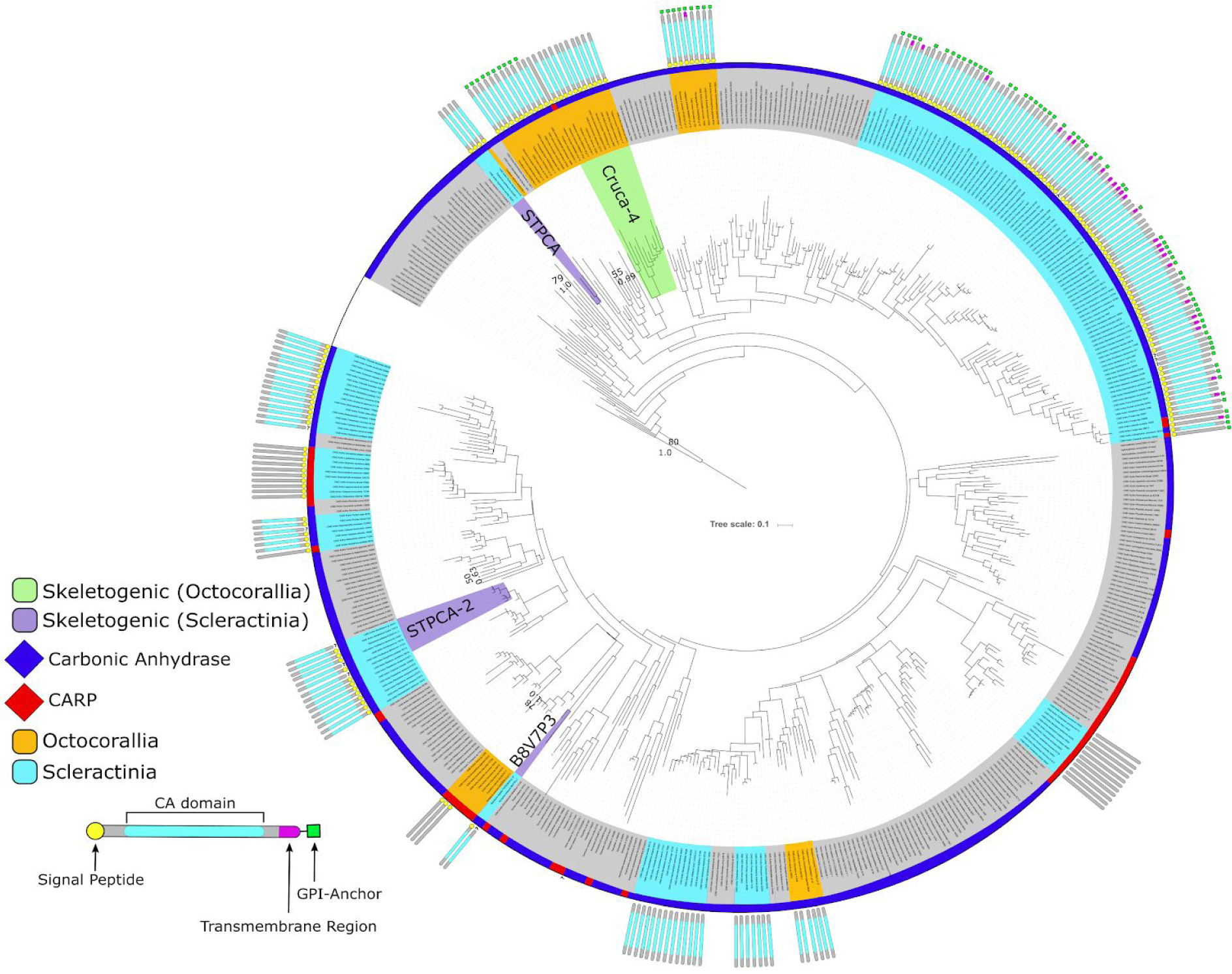
Maximum-likelihood analysis (400 bootstrap replicates) of cnidarian carbonic anhydrases and carbonic anhydrases related proteins (CARPs). Sequences aligned with MAFFT. MUSCLE-based phylogeny in S.Fig 5. Best-fit model: WAG+G+I. Involvement of CruCA4, STPCA, STPCA-2 and B8V7P3 based on Le Goff et al. (2016), Moya et al. (2008), Bertucci et al. (2011) and Ramos-Silva et al. (2013) respectively. Other taxa include: *Homo sapiens*, Porifera, Cubozoa, Hydrozoa, Staurozoa, Scyphozoa, Ceriantharia, Actiniaria, Corallimorpharia. Outgroup: *Chlamydomonas reinhaardtii* (P20507) and *Desmodesmus* sp. (AOL92959.1)

The carbonic anhydrase sequence detected in *M. digitata* grouped together with *A. millepora* B8V7P3, but not with STPCA-2 (Bertucci et al. 2011) which is present in the skeleton of *S. pistillata* (Drake et al. 2013). This suggests that scleractinian families might secrete different carbonic anhydrases as part of their skeleton organic matrix. Finally, the third group comprises scleractinian homologs of *S. pistillata* STPCA (Moya et al. 2008).

## DISCUSSION

Determining which morpho-mineralogical features of coral skeletons are biologically controlled, and which result from environmental effects, remains a key unresolved aspect of coral biomineralization. Here we exploited the co-presence of aragonite and calcite-forming species within Octocorallia, a unique feature among Anthozoa. We provide a first insight into the diversity of proteins occluded within the coral’s CaCO_3_ skeleton, i.e., the skeletome, of species employing different calcification strategies. The identification of several octocoral skeleton organic matrix proteins (SOMPs), in addition to providing new targets for follow up research, also allowed to perform comparative analyses with previously published and the newly characterized *M. digitata* proteome. Our work represents the first examination of the diversity of skeletogenic toolkits across Anthozoa and its relation to the variety of biomineralization strategies displayed by this group.

We have reported low overall proteome overlap, while simultaneously highlighting instances of skeletogenic proteins shared both between and within scleractinians and octocorals. Among these, some proteins are associated with skeleton organic matrices occluded in both aragonitic and calcitic skeletons. Protein presence in the skeleton does not automatically constitute evidence for involvement in biomineralization, as random incorporation within the mineral fraction cannot be excluded. Nevertheless, for different SOMPs present in different groups, proteomic-independent information is available, including, among others, galaxins (Reyes-Bermudez et al. 2009) and carbonic anhydrases (Tambutté et al. 2006; Le Goff et al. 2016). This also applies to aspartic acid-rich proteins, on which extensive research has been conducted (Mass et al. 2013, 2016; Von Euw et al. 2017). Thus the acidic proteins found in *H. coerulea* and *T. musica* could be potential key players in the formation of octocoral skeletons and represent interesting targets for future functional investigations. For other proteins, its presence in the skeleton may not be directly linked to calcification, while still be necessary for the process. For instance, protease inhibitors such as the agrin-like proteins found here in octocoral sclerites are common components of skeleton matrices where they likely prevent matrix degradation caused by different proteases (Marie et al. 2010). Also, the hephaestin-like proteins found here in octocoral sclerites could serve as “deposits” for toxic metals, as proposed for scleractinian skeletons (Ramos-Silva et al. 2013). Alternatively, this could also be linked to ultraviolet radiation absorbance of Fe^3+^ and the capacity of the skeleton to serve as an anti-UV defense structure (Reef et al. 2009). Therefore, although not related to biomineralization, the presence of the same protein in sclerites suggests that octocoral skeletal structures might be employed for functions similar to those found among scleractinians. On the other hand, caution has to be exercised when discussing differences between species due to protein absences as the interpretation of protein absences in proteomic data is affected by multiple factors linked to both sample characteristics and sample processing (Chandramouli & Qian 2009; Michalski et al. 2011; Feist & Hummon 2015) that can reduce the detectability of peptides during mass spectrometry. In light of the above, we focused our results and their interpretation on the presence of coral SOMPs for which proteomic-independent information is available.

Of particular interest, is the secretion of octo-galaxin 1 in the sclerites of octocorals. This represents, to our knowledge, the first report of a galaxin-related protein in anthozoan calcitic skeletons. Phylogenetic analyses of galaxin-related proteins (Bhattacharya et al. 2016) suggest that these proteins are polyphyletic in anthozoans, which in turn suggests that octocoral and scleractinian galaxins represent an instance of convergent evolution. Nevertheless, the fact that only *T. musica*’s octo-galaxin 1 possess the endoprotease target motif characteristic of scleractinian skeletogenic galaxins (Fukuda et al. 2003), and that the observed polyphyly within galaxin phylogenies are likely affected by the inclusion of false homologs, sensitivity to analytical parameters, and aligning algorithms (Conci et al. 2019) makes difficult to rule out the hypothesis of an ancient, biomineralization-related recruitment of (octo)galaxins prior to the divergence of octocorals from the remaining Anthozoa. Similarity in protein features between galaxins *sensu stricto* (see Conci et al. 2019) and galaxin-like proteins, combined with the current lack of support for the deep phylogenetic relationships among these proteins, make understanding the evolution of galaxins difficult and future studies should attempt to provide robust phylogenies for these proteins in order to assess whether their recruitment for biomineralization in Anthozoa is ancient or convergent.

In addition to the presence of homologous skeletogenic components in the skeleton of scleractinians and octocorals, biomineralization in the latter appears to be characterized by evolutionary processes previously proposed for scleractinians, like the enrichment of aspartic acid residues within non-acidic proteins (Takeuchi et al. 2016; Bhattacharya et al. 2016) and the recruitment for calcification of proteins with diverse ancestral biological functions. As for scleractinian galaxin (Forêt et al. 2010), the ubiquitous expression of scleritin homologs across Octocorallia, and its restricted presence in the skeleton of sclerite-forming species suggests a different ancestral function for this protein and a subsequent recruitment for calcification, consistent with the hypothesis of a biomineralization-related recruitment of galaxins for biomineralization in octocorals. Although genomic data remains essential to assess and compare scleritin repertoires in soft corals, the presence of multiple scleritin homologs in *C. rubrum* and *T. musica* reported here points to a gene expansion of scleritins in species forming sclerites. The extent and taxonomic distribution of these expansions, as well as their evolutionary dynamics, remain to be determined once a better sampling of octocoral genomes is available.

While CaCO_3_ polymorph and biomineralization strategy do appear to be correlated with presence/absence patterns of some octocoral skeletogenic proteins, the involvement of CruCA4 in octocoral calcification appears independent of these factors. The current lack of support for deep divergence events during the evolutionary history of Octocorallia does not allow us to provide time estimates for the involvement of CruCA4 in mineralization in octocorals. Efforts to resolve deep divergences in Octocorallia are currently hampered by several factors including rapid radiation (McFadden et al. 2006), slow mitochondrial evolution and inconsistent results between nuclear and mitochondrial markers (see McFadden et al. (2010) for review). The last comprehensive phylogenetic analysis of the subclass split the group into three major clades (McFadden et al. 2006). Species from the genera *Sinularia* and *Tubipora* grouped within the same group, while *Corallium* and *Heliopora* species were included in the other two clades indicating that CruCA4 function as a skeletogenic protein in species belonging to all major octocoral clades and that its involvement in calcification occurred very early on in the evolutionary history of the group.

Finally, within scleractinians, proteomic characterization of the *M. digitata* OM found several putative orthologs of *A. millepora* skeletogenic proteins being secreted into the skeleton. These include acidic proteins from both the SAARP and SAP families and the acidic SOMP B3EWY7. Although the number of shared proteins represent less than a quarter of the overall *A. millepora* skeleton proteome - a sensibly lower percentage than the one between *A. millepora* and *A. digitifera* - these proteins account for nearly 90% of the peptides detected by mass spectrometry in *A. millepora*. This suggests that all major components of the acroporid *toolkit* have been successfully retrieved in *M. digitata*, highlighting a high degree of conservation of the *skeletome* of Acroporidae despite a low fraction of proteins being shared. The discovery and description of skeletogenic proteins represents one of the first essential steps to study biological control in animal biomineralization. Here we have applied a proteomics-based approach to identify and characterize the skeletal proteome of different coral species, covering the diversity of calcification strategies displayed across Anthozoa. In addition to contribute new evolutionary insights on coral biomineralization, this work provides several new targets for future functional investigations. The new data availability for both calcite and aragonite-forming octocoral species is of particular interest, as it opens up the possibility for *in vivo* investigations on biological control over CaCO_3_ polymorph in corals.

## Supporting information

References for sequences datasets used

Homologs distribution for octocoral and scleractinian SOMPs

MSMS_data

Fasta files of the proteomes

## Acknowledgements

SV was supported by the German Research Foundation (DFG) grant Va1146-2/1“MINORCA”. GW was supported by LMU Munich’s Institutional Strategy LMUexcellent within the framework of the German Excellence Initiative, and the German Research Foundation (DFG) grant Wo896/18-1 “MINORCA”, and from the European Union’s Horizon 2020 research and innovation programme under the Marie Skłodowska-Curie grant agreement No 764840 (ITN IGNITE). We thank Dr. Peter Naumann for technical assistance and maintenance of the aquaria facilities and coral culturing, as well as Simone Schätzle and Gabriele Büttner for assistance in the laboratory. SV is indebted to N. Villalobos Trigueros, M. Vargas Villalobos, S. Vargas Villalobos and S. Vargas Villalobos for their constant support.

---

**S.Fig.1.**
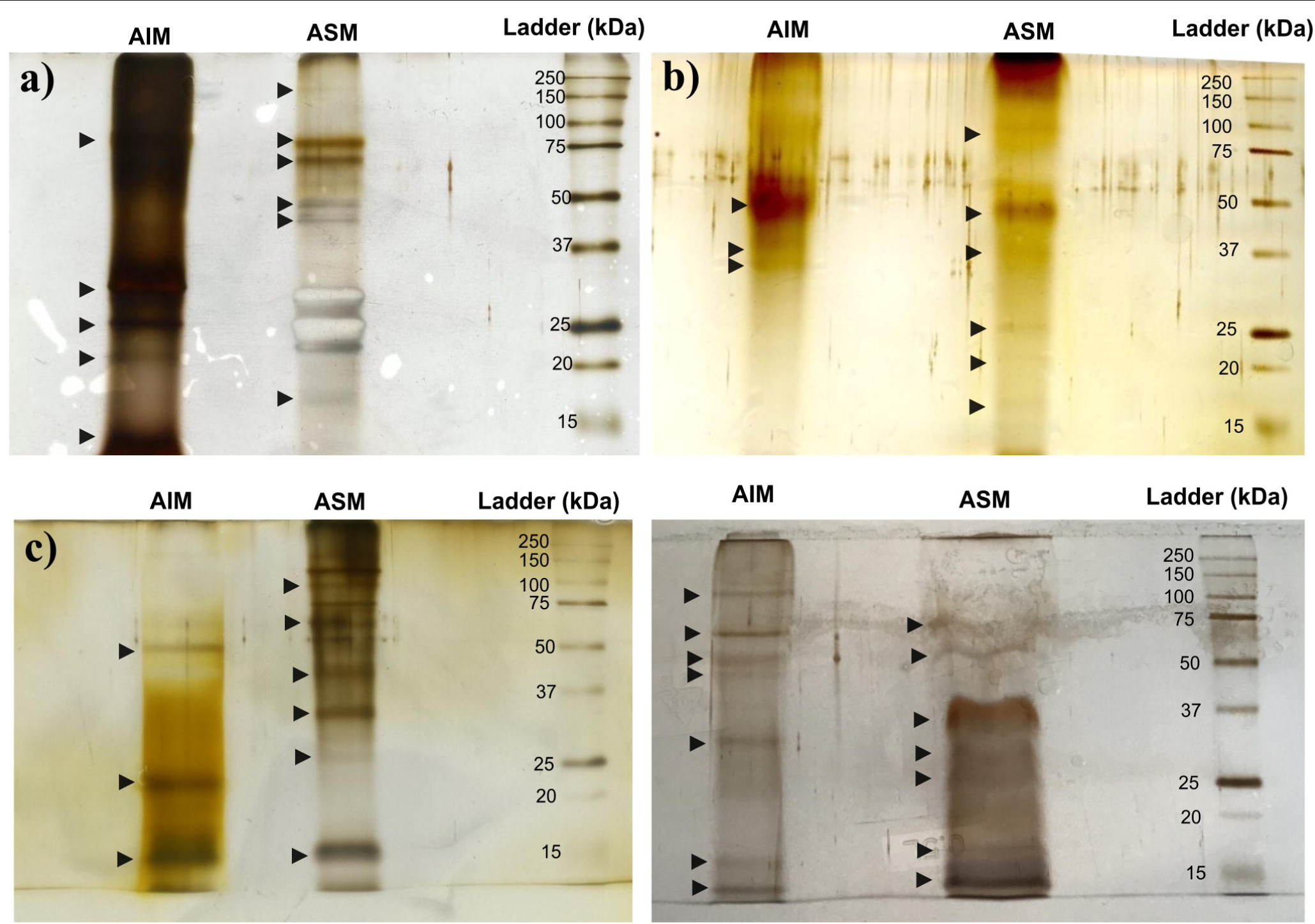
Silver-stained 1D SDS gels of **a)** *H. coerulea*, **b)** *M. digitata*, **c)** *T musica* and **d)** *S. cf. cruciata*.

**S.Fig.2.**
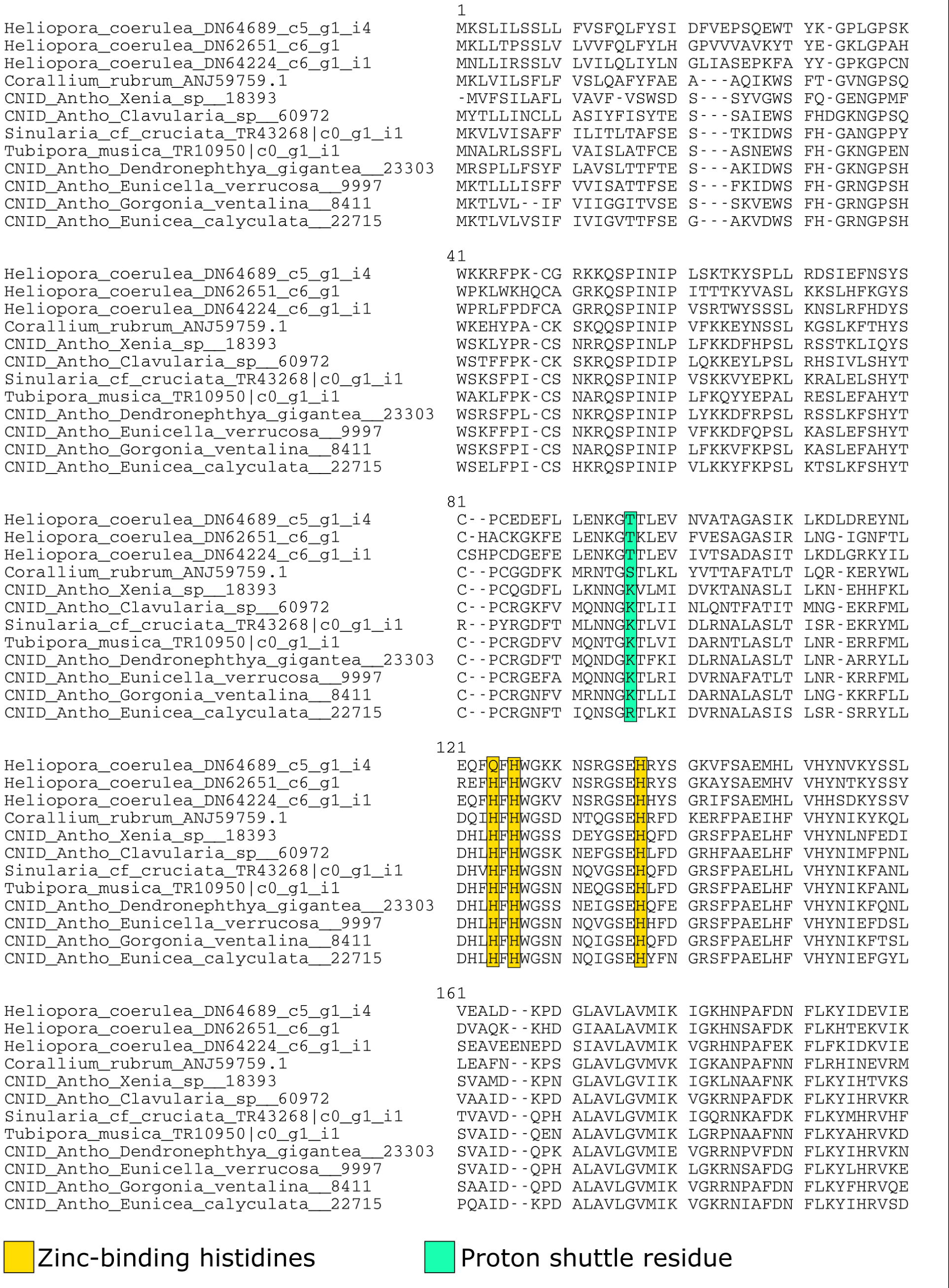
Alignment (MUSCLE) of octocoral CruCA4 homologs. Position of zinc binding and proton shuttle residues based on Del Prete et al. (2017).

**S.fig.3.**
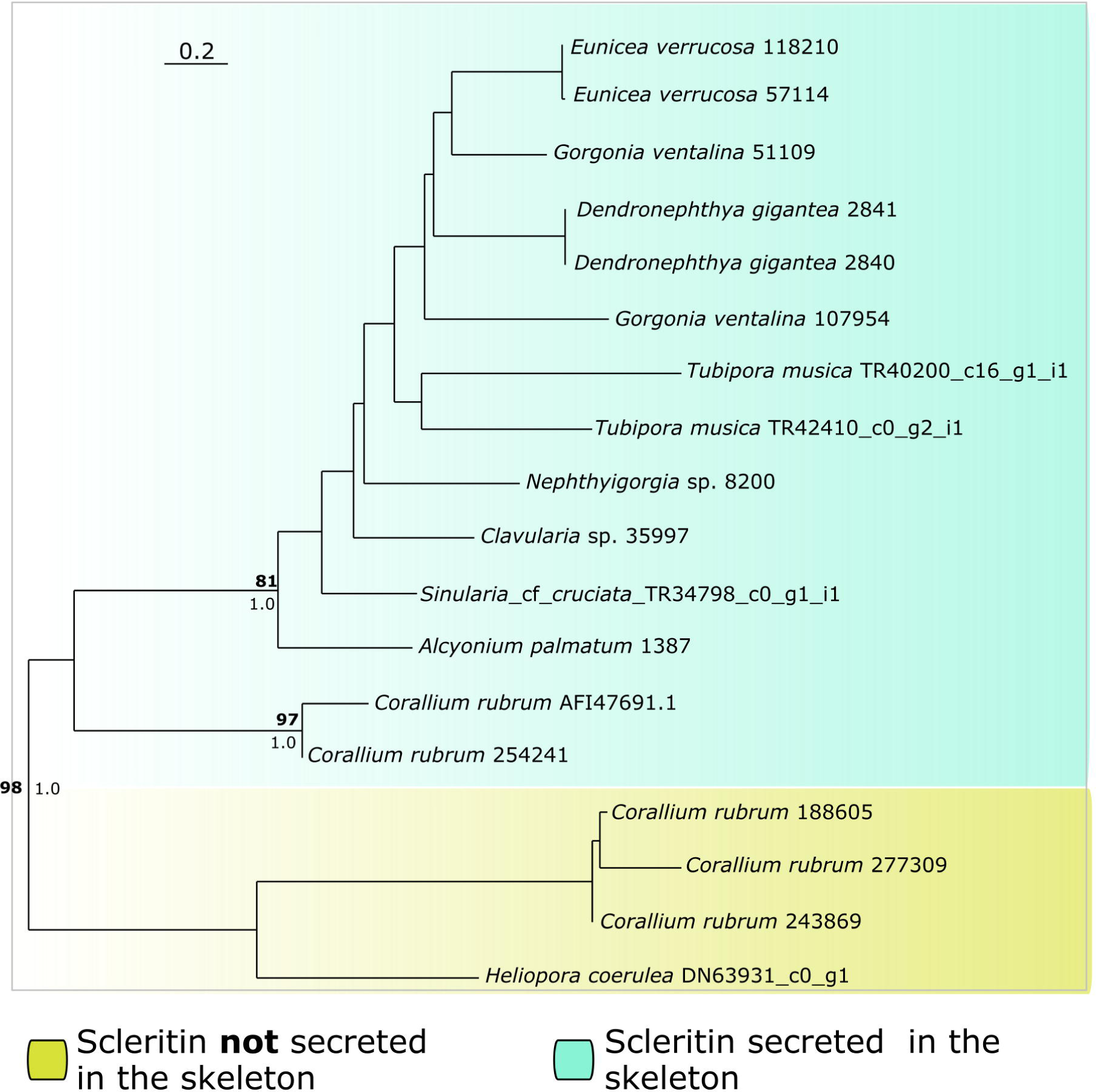
Phylogenetic analysis (400 bootstrap replicates) of octocoral scleritin homologs. Aligning algorithm: MAFFT. Best-fit model: LG+G+ I. Number on nodes: bootstrap support and posterior probability values. Support values in bold represents nodes supported in the MAFFT-based phylogeny. Support showed for nodes of interes only.

**S.fig.4.**
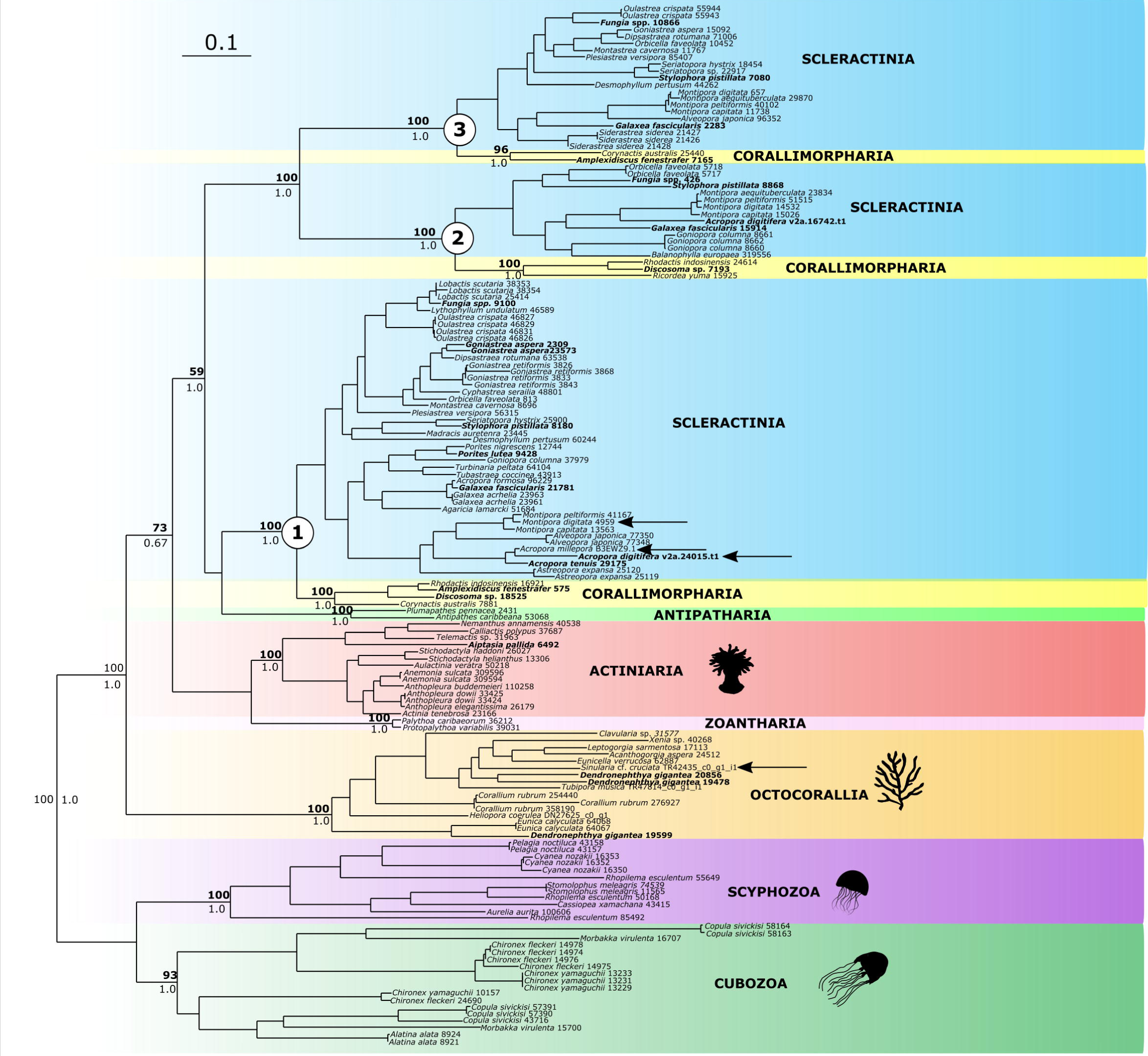
Phylogenetic analysis (400 bootstrap replicates) of cnidarian multicopper oxidases (MCOs). Aligning algorithm: MUSCLE. Best-fit model: WAG+G+I. Number on nodes: bootstrap support and posterior probability values. Support values in bold represents nodes supported in the MAFFT-based phylogeny.

**S.Fig.5.**
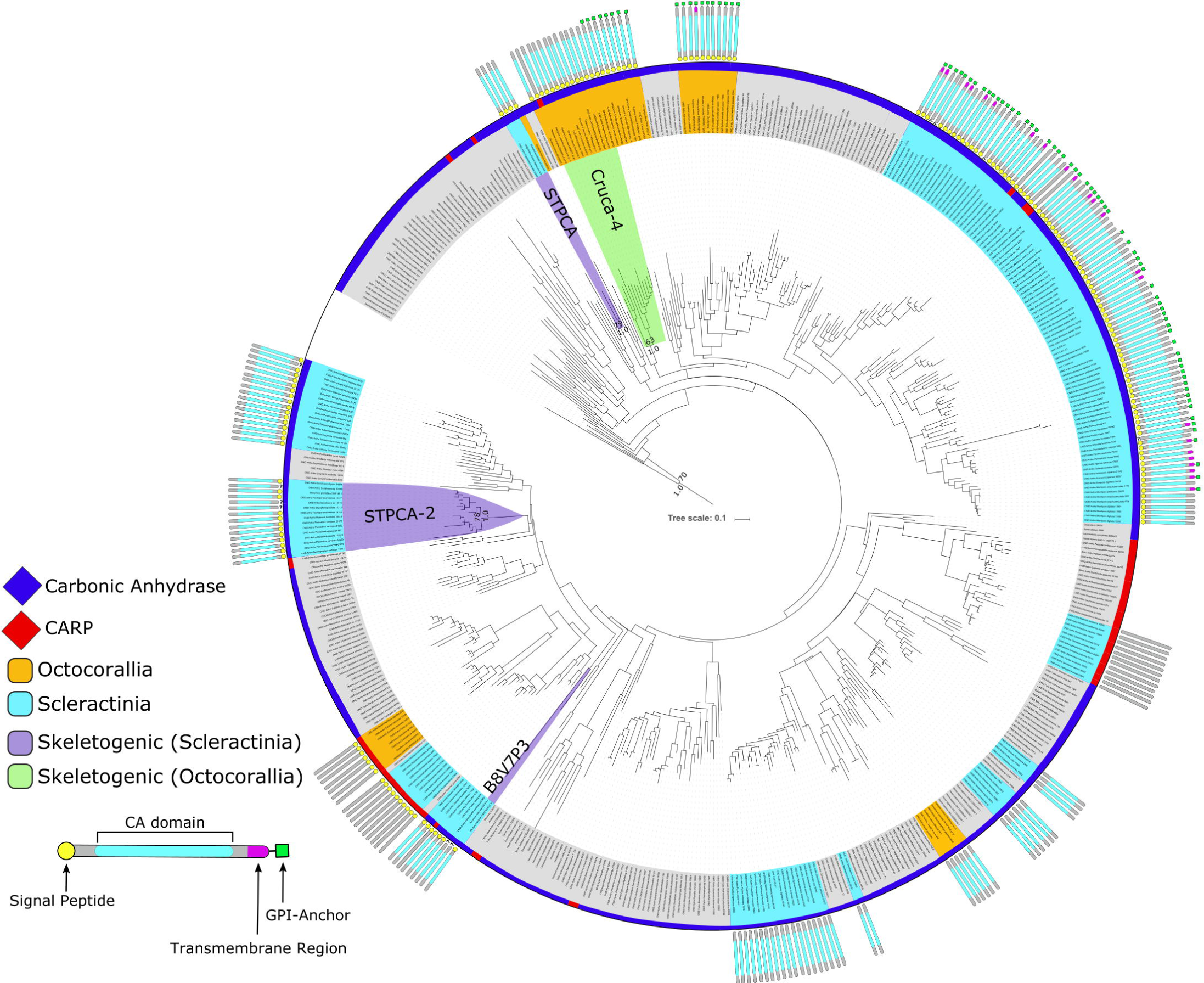
Maximum-likelihood analysis (400 bootstrap replicates) of cnidarian carbonic anhydrases and carbonic anhydrases related proteins (CARPs). Sequences aligned with MUSCLE. Best-fit model: WAG+G+I. Involvement of CruCA4, STPCA, STPCA-2 and B8V7P3 based on Le Goff et al. (2016), Moya et al. (2008), Bertucci et al. (2011) and Ramos-Silva et al. (2013) respectively. Other taxa include: Homo sapiens, Porifera, Cubozoa, Hydrozoa, Staurozoa, Scyphozoa, Ceriantharia, Actiniaria, Corallimorpharia. Outgroup: Chlamydomonas reinhaardtii (P20507) and Desmodesmus sp. (AOL92959.1).

